# A humanized *Caenorhabditis elegans* model of Hereditary Spastic Paraplegia-associated variants in kinesin light chain KLC4

**DOI:** 10.1101/2023.01.07.523106

**Authors:** Selin Gümüşderelioğlu, Lauren Resch, Trisha Brock, Undiagnosed Diseases Network, G.W. Gant Luxton, Queenie K-G Tan, Christopher Hopkins, Daniel A. Starr

## Abstract

Hereditary spastic paraplegia (HSP) is a group of degenerative neurological disorders. We identified a variant in human kinesin light chain *KLC4* that is suspected to be associated with autosomal dominant HSP. How this and other variants relate to pathologies is unknown. We created a humanized *C. elegans* model where *klc-*2 was replaced with human *KLC4* and assessed the extent to which *hKLC4* retained function in the worm. We observed a slight decrease in motility but no nuclear migration defects in the humanized worms, suggesting that *hKLC4* retains much of the function of *klc-2*. Five *hKLC4* variants were introduced into the humanized model. The clinical variant led to early lethality with significant defects in nuclear migration when homozygous, and a weak nuclear migration defect when heterozygous, possibly correlating with the clinical finding of late onset HSP when the proband was heterozygous. Thus, we were able to establish humanized *C. elegans* as an animal model for HSP and use it to test the significance of five variants of uncertain significance in the human gene *KLC4*.

**Summary Statement:** We identified a variant in *KLC4* associated with Hereditary Spastic Paraplegia. The variant had physiological relevance in a humanized *C. elegans* model where we replaced *klc-2* with human *KLC4*.

## Introduction

Over 10,000 disorders are classified as rare diseases, each affecting fewer than 1/2000 people (Ferreira, 2019). Together, they are not rare; over 4% of the world’s population is currently suffering from a rare disease (Nguengang Wakap *et al*., 2020). Diagnoses, let alone treatments, of rare diseases are difficult because underlying mutations are spread over 8,000 genes (Ferreira, 2019). Even whole-genome sequencing leads to a definitive diagnosis only about 25% of the time (Smedley *et al*., 2021). More often, a definitive diagnosis is not returned, and the clinician is left with a list of variants of uncertain significance and little idea as to which of these variants are pathogenic. Thus, one of the biggest challenges in genomic medicine is the validation of which identified variant is pathogenic. The bottleneck facing clinical geneticists is a need for functional data that can assess the pathogenicity of a variant of uncertain significance.

Hereditary Spastic Paraplegia (HSP) is a group of monogenetic diseases that are classified as rare diseases that present at various times throughout life. Individuals characteristically suffer from neurodegeneration in the longest motor neurons, leading to progressive spasticity and lower limb weakness (Parodi *et al*., 2017; Gumeni *et al*., 2021; Shribman *et al*., 2019). Upwards of 79 genes have been linked to HSP, yet geneticists fail to obtain definitive genetic diagnoses in over half of suspected HSP individuals (Parodi *et al*., 2017; Gumeni *et al*., 2021; Shribman *et al*., 2019). This suggests that mutations in additional unknown genes lead to HSP. Moreover, once new candidate HSP genes are identified, we need an *in vivo* model to access the physiological significance of newly identified variants for a timely clinical diagnosis (Hopkins *et al*., 2022).

Functional studies *in vivo* are important for variant assessment. *Caenorhabditis elegans* is a model system that can relatively inexpensively test variants of uncertain significance at the speed needed for inclusion in a clinical report (Baldridge *et al*., 2021). *C. elegans* also allows examination of function in the context of a developing tissue and the use of a variety of biochemical, developmental, and quantitative cellular assays needed to detect subtleties of variant biology. These advantages have led to many reports modeling human diseases in *C. elegans* (Kropp *et al*., 2021). Thus, humanized *C. elegans* models are likely to be useful in testing the *in vivo* consequences of variants of uncertain significance identified in the clinic. Here, we report the design and use of a humanized *C. elegans* model to test the clinical significance of variants of uncertain significance in the human kinesin light chain gene *KLC4* identified in individuals with HSP.

Molecular motor-based transport along microtubules is essential for the function and survival of eukaryotic cells (Ross *et al*., 2008). Microtubule motors are especially important in transporting organelles and molecules down the length of long motor neuron axons (Hirokawa *et al*., 2010; Perlson *et al*., 2010; Saxton and Hollenbeck, 2012). Disrupting motors leads to a variety of neurodegenerative diseases (Mandelkow and Mandelkow, 2002; Kurd and Saxton, 1996; Giudice *et al*., 2014). Kinesin-1 is the founding member of the kinesin superfamily of microtubule motors (Vale *et al*., 1985). Kinesin-1 consists of a tetramer of 2 kinesin light chains that bind to the tails of 2 kinesin heavy chains. The heavy chains, called Kif5b in humans, bind microtubules and provide the ATPase motor activity while the light chains serve as cargo adapters (Verhey *et al*., 1998). In the presence of a cargo bound to the light chains, kinesin-1 is activated to move towards the plus end of microtubules. In humans, there are four different kinesin-1 light chains; KLC1, KLC2, KLC3 and KLC4. While KLC1, KLC2 and KLC3 are relatively well studied (Rahman *et al*., 1998; Junco *et al*., 2001; Zhang *et al*., 2012) and their functions as well as how they are involved in certain diseases are known, KLC4 is under studied. Yet, mutations in *KLC4* are linked to diseases including lung cancer (Baek *et al*., 2018; 2020) and HSP (Bayrakli *et al*., 2015). The goal of this project is to further explore the link between *KLC4* and HSP by developing a humanized *C. elegans* model as a clinical avatar to test the functions of variants of uncertain significance in the gene *KLC4*.

## Materials and Methods

### *C.elegans* genetics and humanized strain generation

*C. elegans* strains were maintained on nematode growth medium plates seeded with OP50 *E. coli* at room temperature; the N2 strain was used as wild type (Brenner, 1974). Some strains were obtained from the Caenorhabditis Genetics Center, funded by the National Institutes of Health Office of Research Infrastructure Programs (P40 OD010440). The *ycIs9 I* strain, which was used to mark hypodermal nuclei with GFP, was generated along with *ycIs10 V* as previously described (Bone *et al*., 2014). The strains used in this study are listed in Table 1.

**Table 1.**
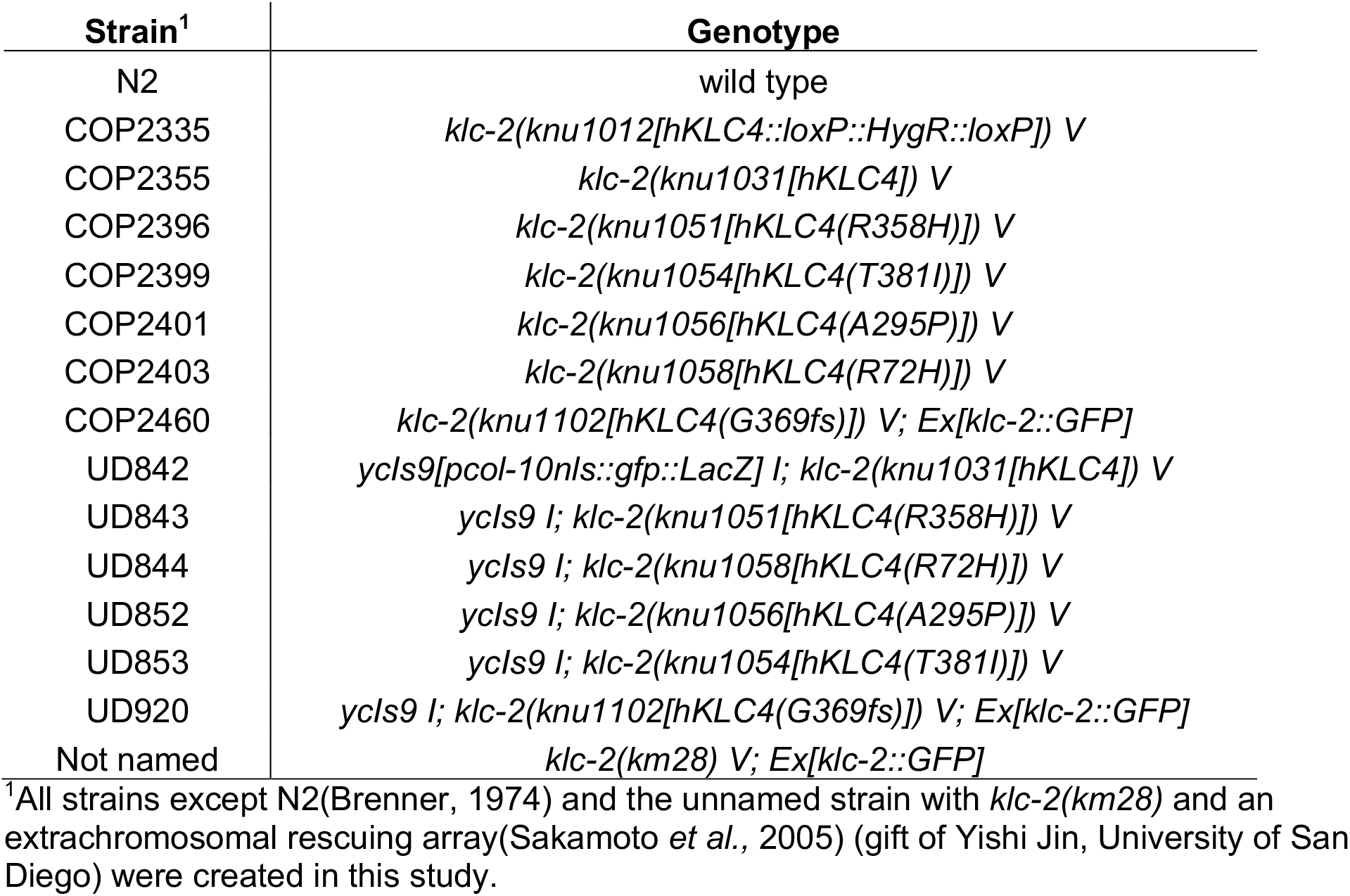
*C. elegans* strains used in this study.

The *hKLC4* strain was made utilizing the transgenesis services of InVivo Biosystems. To generate the *hKLC4* strain, the coding region of the human *KLC4* open reading frame was synthesized from the ATG to the stop codon of the most supported (Transcription Support Level 1) KLC4 isoform in Ensembl (ENST00000347162.10) and placed into a plasmid (Guo *et al*., 2014). The resulting gene coding for 619 amino acids was codon-optimized for *C. elegans* (Mitreva *et al*., 2006), and synthetic introns were inserted, which has been shown to be essential for normal expression in *C. elegans* (Blumenthal, 2012). The sequence of this gene block is shown in Supplemental Material. The *hKLC4* coding sequence was then flanked by 500 bp endogenous *C. elegans klc-2* 5’ and 3’sequences from genomic DNA by PCR and Gibson assembly to create pNU2756. A hygromycin resistant gene with a *tbb-2* 3’UTR selection cassette flanked by *loxP* sites was included in pNU2756 to aid in identifying transgenic animals. We also included the 3’UTR sequence of *eft-3* after of the *hKLC4* stop codon because sometimes a strong 3 ‘UTR is needed for optimal expression (Chen *et al*., 2013). pNU2756 was used as the repair template for CRISPR/Cas9 genome editing (Figure 1). Two sgRNAs targeting each end of the *klc-2* open reading frame were used to guide CRISPR/Cas9 to cut out the coding region of *klc-2* (Table 2). The sgRNAs that were preassembled with crRNA, Cas9 protein and the assembled ribonucleoprotein complex, and the repair template were then injected into the gonads of *C. elegans* young adults (Paix *et al*., 2015; Farboud *et al*., 2019). Progeny were screened for incorporation of *hKLC4* into the worm genome by selecting for the animals that could survive upon hygromycin treatment. Two strains with *hKLC4::eft-3 3’UTR* and the hygromycin selection cassette were obtained and backcrossed to N2 wild type to minimize off-target effects. Expression of *hKLC4* was confirmed by RT-qPCR. Finally, using new sgRNAs (Table 2), the *hygromycin::tbb-2 3’UTR* cassette and the *eft-3 3’UTR* were removed, restoring the native *klc-2 3’UTR* and generating the humanized *klc-2(knu1031[hkLC4])* strain, hereafter referred to as *hKLC4* (Figure 1).

**Table 2.**
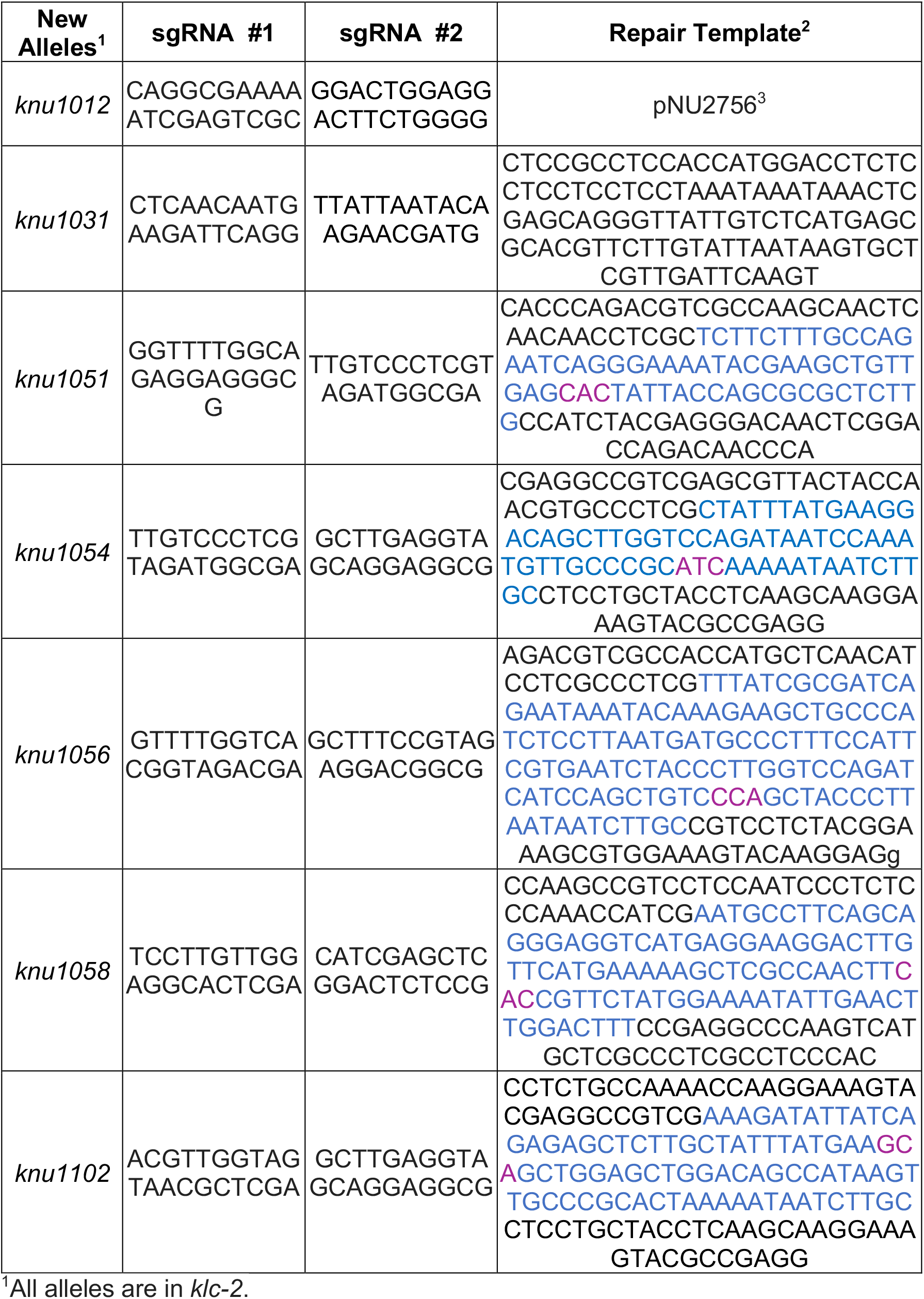
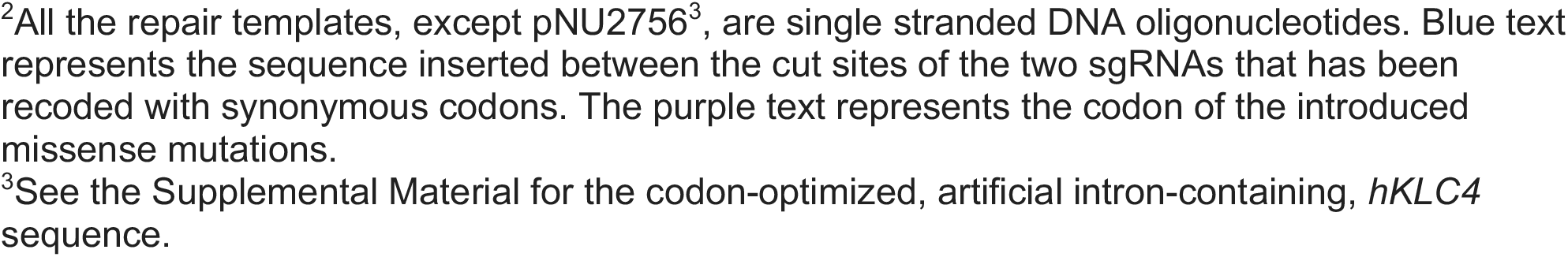
sgRNA and repair DNA templates used in this study.

**Figure 1:**
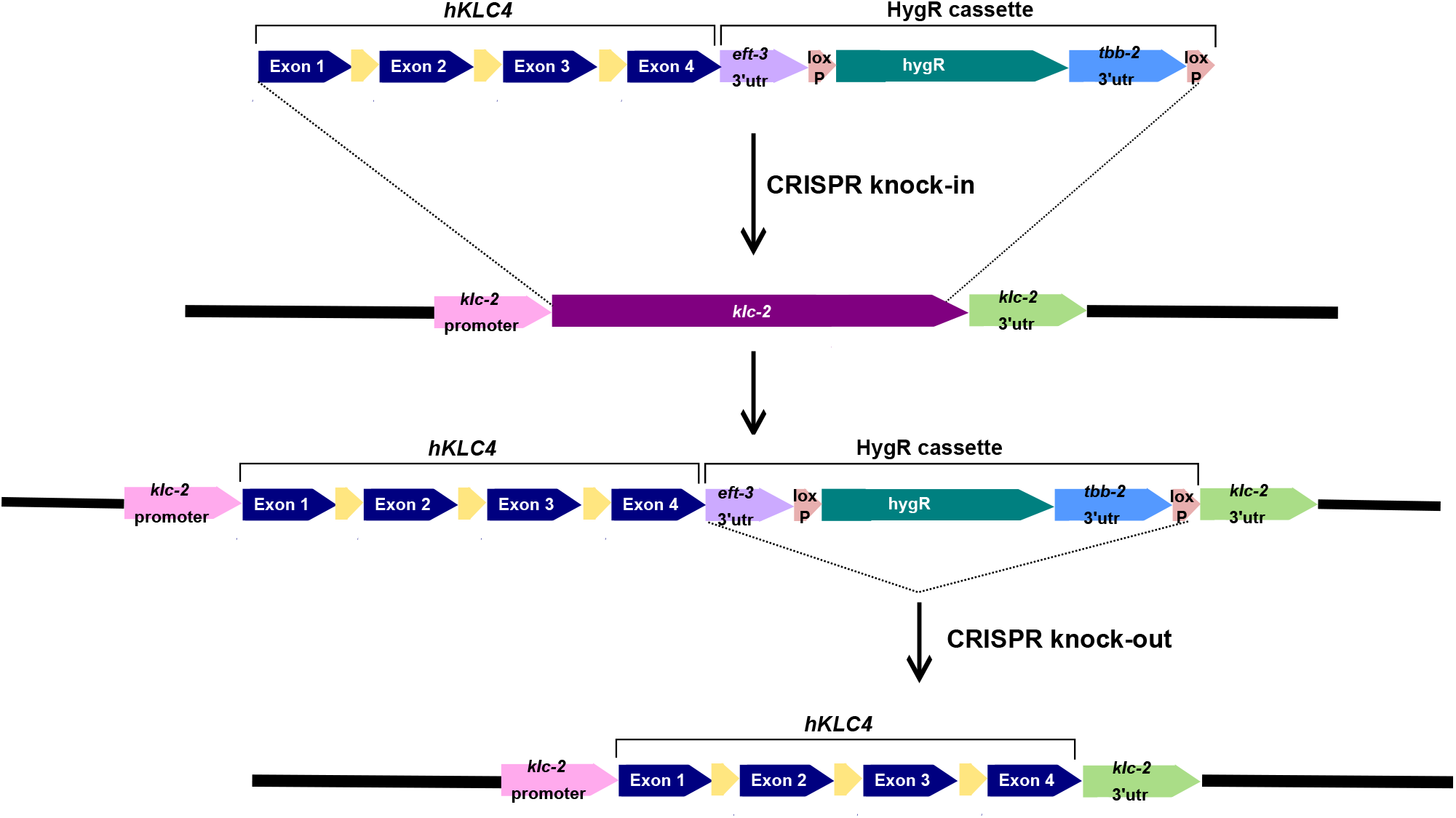
The CRISPR/Cas9-mediated genome editing workflow used to generate *hKLC4* worms. Inserted sequence contains human gene *KLC4* (exons shown in blue; synthetic introns shown in yellow) and hygromycin gene resistant selection cassette (HygR). See text for details.

Point mutations were introduced into the *hKLC4* line using CRISPR/Cas9 gene editing (Farboud *et al*., 2019). The sgRNA and ssDNA repair template sequences used to introduce missense mutations studied in this work are listed in Table 2. To identify successfully injected animals, co-CRISPR with templates to create *dpy-10(gof)* alleles was performed (Arribere *et al*., 2014). Point mutants were then screened for with genomic PCR followed by restriction digest analysis. Newly generated alleles were backcrossed to N2.

To generate the humanized allele with the clinical variant of uncertain significance, *klc-2(knu1102[hKLC4(G369fs)])*, wild-type *klc-2::gfp* expressed from an extrachromosomal array (Sakamoto *et al*., 2005) was crossed into the *hKLC4* strain. Then, CRISPR/Cas9 gene editing was used as described above to introduce the mutation into *hKLC4*. By having a *klc-2::gfp* array, the worms were able to mimic a heterozygous condition for this variant, which was similar to the individual who was also heterozygous. To further analyze the clinical variant for haploinsufficiency, heterozygous *hKLC4* G369fs/+ animals were generated by mating *hKLC4* males with *hKLC4* G369fs, *klc-2::gfp* hermaphrodites, and selecting for animals without the *klc-2::gfp* extrachromosomal array.

### Phenotypic assays and statistical evaluations

To quantify nuclear migration, humanized *hKLC4* strains were crossed into a GFP nuclear marker expressed in larval hypodermal nuclei (*ycIs9 I*; Table 1). Nuclear migration assays were performed as described (Fridolfsson *et al*., 2018). Briefly, L1-2 worms with GFP-marked hypodermal nuclei were picked and mounted on 2% agarose pads in ∼5 µl of 1 mM tetramisole in M9 buffer. Syncytial hyp7 nuclei were scored as abnormally located if they were in the dorsal cord.

For brood size assays, starting at the L4 stage, 10-15 single animals of each genotype were transferred onto fresh OP50 *E. coli* plates, labeled as Day 1, and kept at room temperature (approximately 22°C) for 42 hours so that they became adults and had about 24 hours to lay eggs. At 42 hours, adult worms were moved to new plates, labeled as Day 2, and kept at room temperature for 24 hours. At the 24-hour mark adult worms were moved to new plates, labeled Day 3, and dead eggs and young worms from the Day 1 plates were counted. At the next 24-hour mark, the adult worms on the Day 3 plates were killed and dead eggs and young worms from Day 2 plates were counted. At the next and final 24-hour mark, dead eggs and young worms from Day 3 plates were counted.

For the motility assays, 8-10 L4 stage animals were put into a plate and flooded with M9 buffer. We observed and filmed worms swimming in buffer for 30 seconds. Using the Fiji wrMTrck plugin (Nussbaum-Krammer *et al*., 2015), we measured the number of body bends per second (BBPS).

Prism nine software was used for the statistical analyses. All the data from nuclear migration, brood size and lethality, and motility assays were displayed as scatter plots with means and 95% confidence interval (CI) as error bars. Sample sizes and the statistical tests are indicated in the figure legends. Unpaired student t-tests were performed on the indicated comparisons.

## Results

### Clinical description of an affected individual with HSP

A male individual with the clinical *KLC4* variant presented to the Undiagnosed Diseases Network with slowly progressive myelopathy, radiculopathy, and neuropathy since around 50 years of age. Initial symptoms included numbness, proceeded by weakness and lower extremity hypertonia, hyperreflexia and spasticity. The individual’s symptoms worsened over the next twenty years, although he maintained normal cognitive ability. Ophthalmologic evaluation showed thinning of the ganglion cell layer and papillomacular bundle, but no visual changes. Of note, he also had celiac disease, which can cause myelopathy and neuropathy, but he had been compliant with treatment with no obvious symptoms. The individual has a healthy sibling who did not harbor the *KLC4* variant. The individual worked as an agronomist and was frequently exposed to herbicides and pesticides, potentially complicating the diagnosis.

### Selection of variants of *KLC4* to aid molecular diagnosis of individuals with HSP

A pathogenic variant was identified in the individual described above with HSP where a GG pair of nucleotides in the open reading frame of *KLC4* was deleted to cause a frame shift (NM_201523.2; c.1160-1161delGG; p.G369Afs*8) (Supplemental Figure 1). The predicted mutant KLC4 protein (G369fs) replaces the glycine at position 369 with an alanine, followed by eight additional novel residues and a premature stop codon that truncates more than a third of the protein. The probability of loss of function (pLOF) score for *KCL4* is 0.53 (database https://gnomad.broadinstitute.org/ - assessed July 24, 2022), an intermediate value for essentiality, suggesting that in a subset of genetic contexts, the gene variants can be associated with an autosomal dominant disorder. There was no evidence of any duplication or deletion of the *KLC4* gene.

We turned to bioinformatic databases in attempts to identify two predicted pathogenic and two predicted benign variants as reference alleles in addition to the clinical *KLC4* G369fs variant (Supplemental Figure 1). Two missense *KLC4* mutations, R72H and R358H, were chosen as predicted benign controls. R72H was observed at a very high frequency (6803x) in healthy populations using GnomAD (Karczewski *et al*., 2020). R358H was seen at 9x in GnomAD and was scored as possibly damaging in PolyPhen-236, tolerated in SIFT37, and neutral in CADD (Kircher *et al*., 2014) and REVEL (Ioannidis *et al*., 2016).Two other missense mutations were chosen because they were predicted to be pathogenic variants. Both T381I and A295P were identified as damaging by PolyPhen-2 (AdzhubeIvan *et al*., 2010) and possibly damaging by SIFT (Sim *et al*., 2012). Thus, we have a collection of five *KLC4* mutations for testing in an *in vivo* model.

### A *C. Elegans* model where human *KLC4* rescues the lethality of a *klc-2* null allele

We aimed to make a humanized *C. elegans* model to test the physiological significance of *KLC4* mutations. Kinesin-1 plays similar important roles in *C. elegans* as it does in humans, including moving synaptic vesicles in motor neurons and nuclei in hypodermal precursors (Meyerzon *et al*., 2009; Sakamoto *et al*., 2005). *klc-2* encodes what is likely the primary light chain for kinesin-1 in *C. elegans*. Null alleles of *klc-2* cause larval lethality while *klc-1* is divergent, not essential, and does not bind kinesin heavy chain (UNC-116) (Sakamoto *et al*., 2005). Human KLC4 and *C. elegans* KLC-2 proteins both have a predicted coiled-coil domain that binds to the kinesin heavy chain, and six tetratricopeptide (TPR) repeats that function together to bind cargo (Figure 2A-B). The coiled-coil regions of KLC4 and KLC-2 are 46% identical while the TPR domains are 78% identical (Figure 2A and Supplemental Figure 1). Using AlphaFold (Jumper *et al*., 2021), we were able to model the predicted structure of KLC4 as well as its interactions with both UNC-116 and the *C. elegans* protein UNC-83 that acts as a binding adaptor for kinesin-1 (Meyerzon *et al*., 2009; Taiber *et al*., 2022). Thus, we chose the *klc-2* locus to engineer in human *KLC4* to make a humanized model. Human KLC4 has a major isoform encoding a 619 residue protein that is expressed more broadly and at higher levels than the 637 residue isoform according to the Transcript Support Level (TSL) method (Yates *et al*., 2016). Therefore, we used the shorter 619 protein isoform to make a humanized *C. elegans* line. A humanized *C. elegans* model was generated where the endogenous *klc-2* gene was replaced by human *KLC4* using CRISPR/Cas9 genome editing (Figure 1). The coding region for human *KLC4* was codon optimized, placed under control of the endogenous *klc-2* promoter and the 5’- and 3’-untranslated regions of the *klc-2* gene. In addition, three synthetic *C. elegans* introns were inserted into *KLC4* to maximize its expression in *C. elegans* (Figure 1).

**Figure 2:**
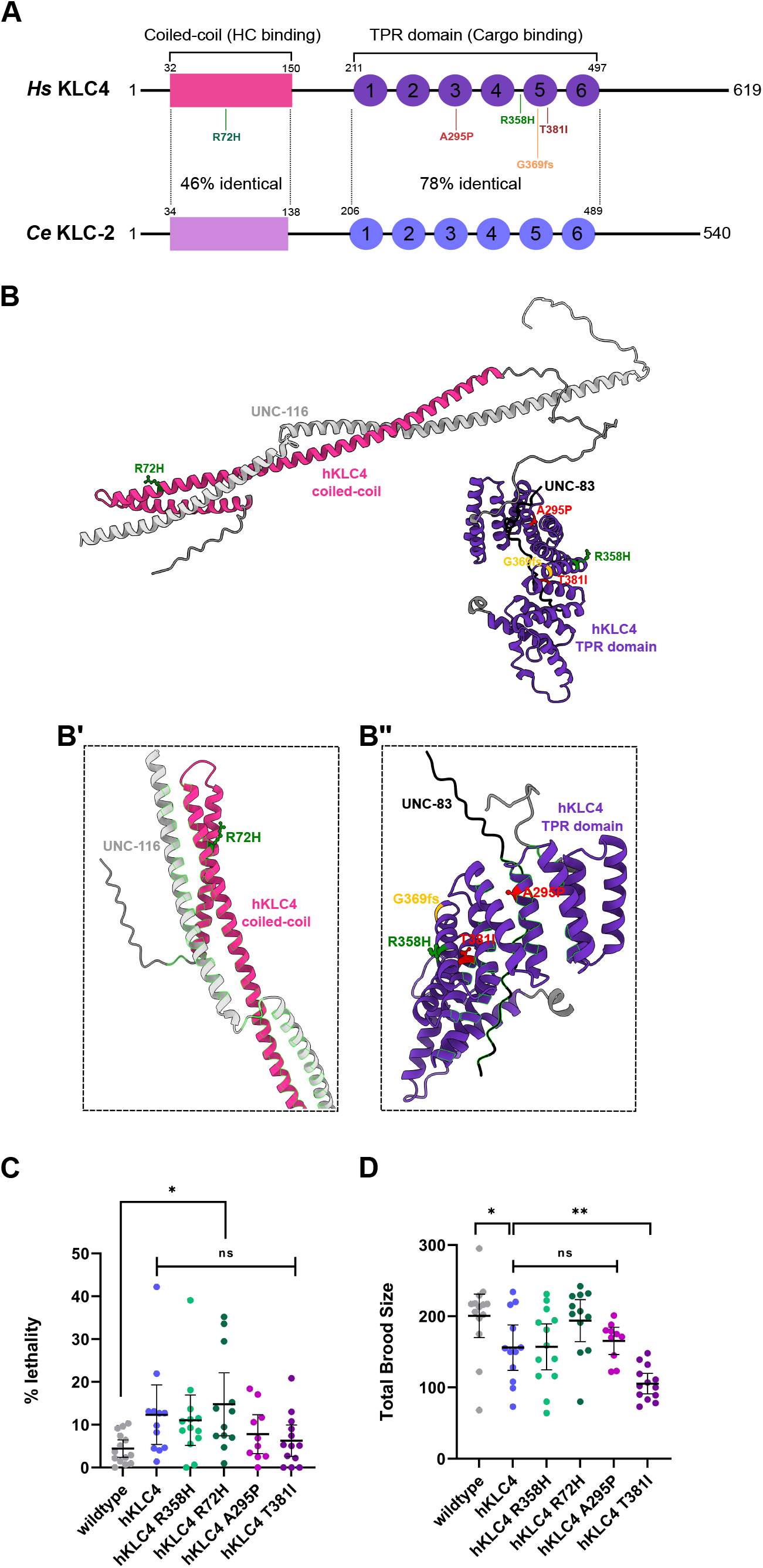
A *C. elegans* model where human *KLC4* replaces *klc-2*. A) Illustration of the human KLC4 and *C. elegans* KLC-2 proteins. The predicted coiled-coil domain (pink) that binds to the kinesin heavy chain, and six TPR repeats (purple) that function together to bind cargo are shown for both proteins. The coiled-coil regions of KLC4 and KLC-2 are 46% identical while the TPR domains are 78% identical. The missense mutations used in this study are shown. Green mutants are predicted benign, red are predicted pathogenic, and the orange mutant is the clinical variant of uncertain significance. B) AlphaFold prediction of the structures and interaction between hKLC4 (coiled-coil domain in pink and TPR domain in purple), the kinesin heavy chain UNC-116 (silver) and UNC-83 (black). The missense mutations used in this study are shown. Green mutants are predicted benign, red mutants are predicted pathogenic, and the orange mutant is the clinical variant of uncertain significance. B’ and B’’ show the predicted interactions between UNC-116 and hKLC4, and between KLC4 and UNC-83, respectively. Interacting residues are highlighted. C) Quantification of the % lethality of *C*.*elegans* strains. D) Quantification of total brood size of *C. elegans* strains. For C-D, each data point represents one animal. n= 10-15 for each strain. Means with 95% CI are shown in error bars. Unpaired student t-tests were performed on the indicated comparisons; ns means not significant, p>0.05; * p<0.05; ** p<0.005.

After generating the humanized *C. elegans* line where human *KLC4* replaced *klc-*2 (hereafter referred as the *hKLC4* line) we compared the fitness of the new model to wild type. We quantified the viability of the *hKLC4* line by measuring its lethality and brood size in comparison to wild type (Figure 2C-D). The *hKLC4* strain was viable as a homozygous strain, while *klc-2* null animals are 100% embryonic or L1 larval lethal, suggesting that *hKLC4* rescued many of the essential *klc-2* functions. However, the *hKLC4* line had significant levels of embryonic lethality (11.2 ± 5.4 % compared to 4.1 ± 1.8% in wild type; mean ± 95% confidence intervals) and a slightly lower brood size (142.6 ± 30.2 compared to 191.1 ± 31.9 in wild type) (Figure 2C-D), suggesting that *hKLC4* animals were not as fit as wild type. Nonetheless, most animals with only human KLC4 in place of endogenous KLC-2 were quite viable, fertile, and appeared relatively normal, suggesting that the *hKLC4* model would be of use as a clinical avatar.

The *hKLC4* animals had no obvious phenotypes affecting their ability to crawl on the surface of an agar plate. However, swimming and crawling are two different forms of movement in terms of their kinematics and muscle activity (Pierce-Shimomura *et al*., 2008). Therefore, to conduct a more comprehensive movement analysis of the humanized animals, we used a swimming assay where we observed head-thrashing in liquid. We scored swimming by counting the number of body bends per second (bbps) to assess the motility of *hKLC4* animals (Pierce-Shimomura *et al*., 2008; Mattout *et al*., 2011). We observed that *hKLC4* animals had a significantly lower number of body bends per second than wild-type worms (0.72 ± 0.21 bbps compared to 1.48± 0.09 bbps in wild type) (Figure 3A, B, C; Videos 1-2). This two-fold effect suggests that the humanized line has a significant motility defect. However, these animals retained most of their swimming activity, suggesting that swimming can serve as a sensitized assay for measuring the effect of variants of uncertain significance.

**Figure 3:**
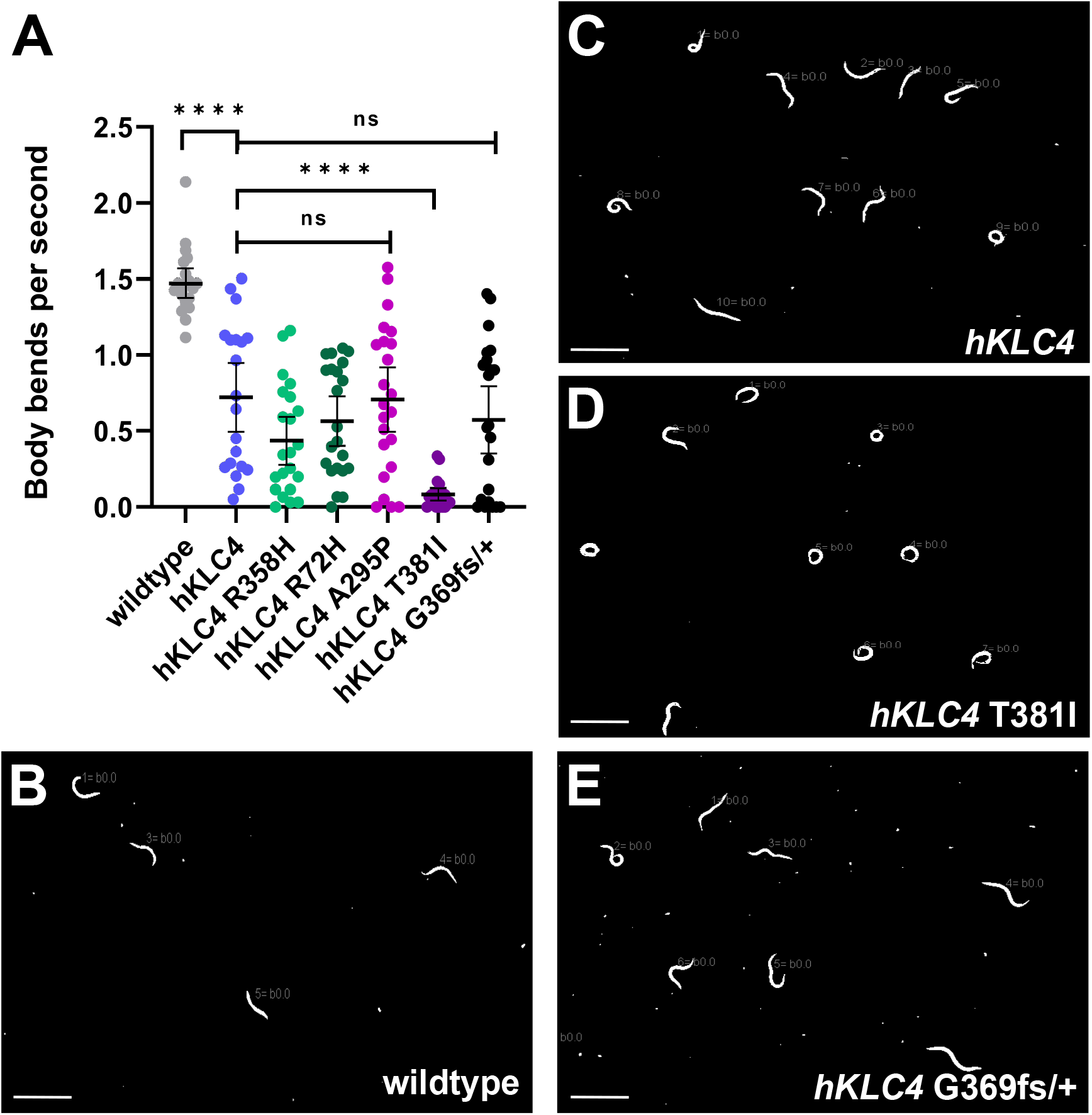
*hKLC4* worms have a motility defect that is enhanced by the predicted pathogenic mutation T381I. A) Quantification of *C. elegans* swimming by counting the number of body bends per second. Each point represents one L4-stage animal. n=20 for each strain. Means with 95% CI are shown in error bars. Unpaired student t-tests were performed on the indicated comparisons; ns means not significant, p>0.05; **** p<0.0001. B-E) Images of (B) wildtype (N2), (C) *hKLC4*, (D) *hKLC4* T381I, and (E) *hKLC4* G369fs/+ animals swimming in buffer. Scale bars, 1mm.

A second *klc-2* dependent assay we examined was nuclear migration (Fridolfsson *et al*., 2018). In mid-embryogenesis, two rows of hyp7 precursors on the dorsal surface of embryos intercalate to form a single row spanning the dorsal midline. Next, nuclei migrate contralaterally toward the plus ends of microtubules across the dorsal midline to the opposite side of the embryo (Sulston *et al*., 1983) (Figure 4A). Successful completion of nuclear migration in embryonic hyp7 hypodermal cells requires kinesin-1 heavy chain and KLC-2 (Meyerzon *et al*., 2009). The linker of nucleoskeleton and cytoskeleton (LINC) complex, consisting of the Klarsicht/ANC-1/SYNE homology (KASH) protein UNC-83 at the outer nuclear membrane and the Sad1/UNC-84 (SUN) protein UNC-84 at the inner nuclear membrane, recruits kinesin-1 to the surface of nuclei and transmits the forces to inside the nucleus (Starr, 2019) (Figure 4A). Null mutant *klc-2(km28)* larvae that barely escape embryonic lethality had an average of 10.4 ± 1.4 hyp7 nuclei abnormally located in the dorsal cord, compared to 0.07 ± 0.08 in wild type (Figure 4B). We have previously shown that this represents a nearly penetrant nuclear migration defect (Fridolfsson and Starr, 2010). To analyze if KLC4 was able to retain KLC-2 function in the humanized worms, we counted the number of hyp7 nuclei present in the dorsal cord. The *hKLC4* animals had no significant nuclear migration defects when compared to the wild type (Figure 4B-C). Together, these data suggest that *hKLC4* can substitute for most of the function of *klc-2* in *C. elegans*.

**Figure 4:**
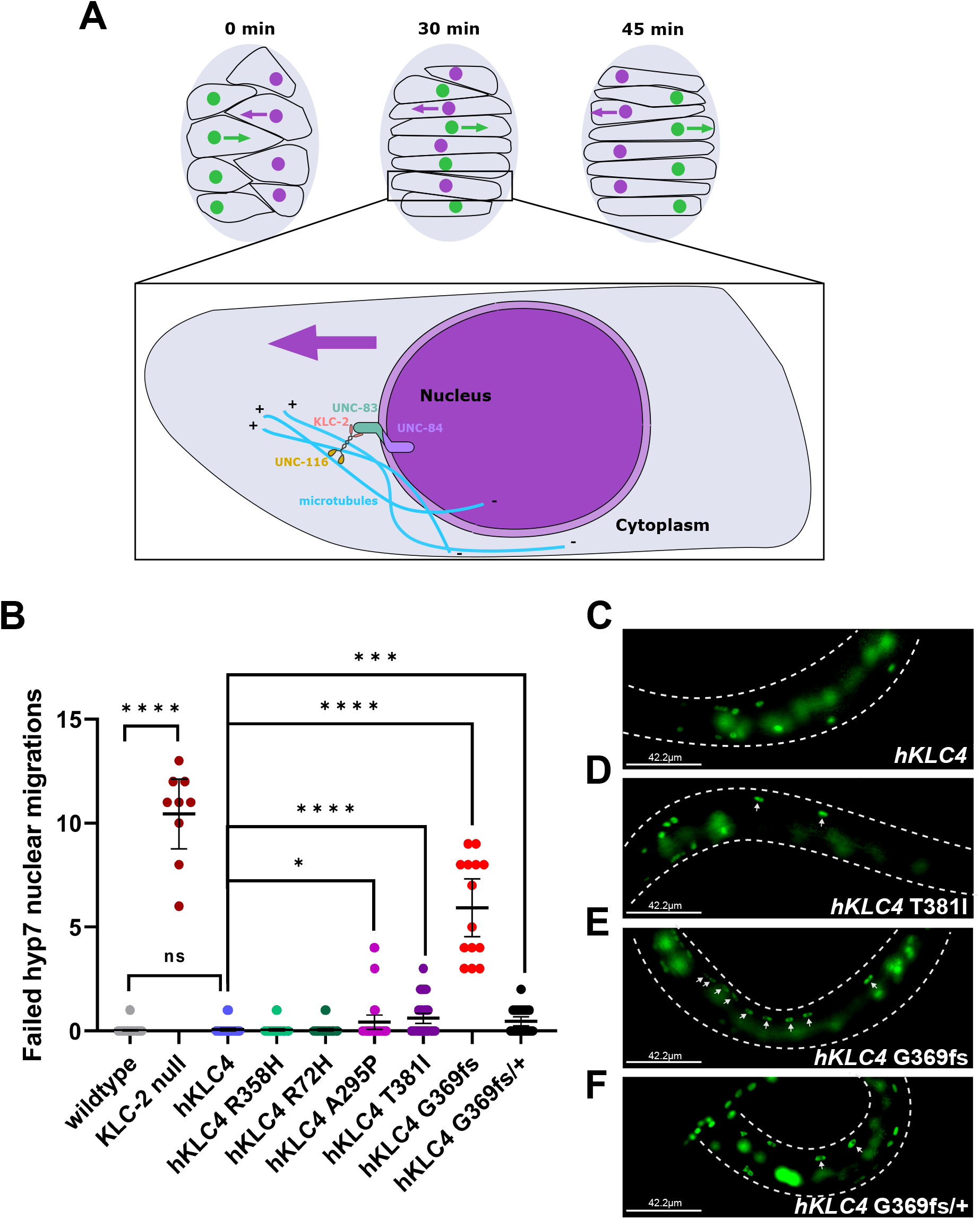
The clinical variant of uncertain significance *hKLC4* G369fs causes a severe hyp7 nuclear migration defect. (A) Illustration of the dorsal view of hyp7 nuclear migration during mid-embryogenesis. At t=0 min, nuclei of hyp7 precursors (green and purple) are found on the right and left sides of the dorsal surface of embryos. At t=30 min, the nuclei, mediated by the LINC complex (UNC-83 and UNC-84), intercalate to form a single row spanning the dorsal midline. At t=45 min, nuclei migrate contralaterally toward the plus ends of microtubules (blue) across the dorsal midline to the opposite side of the embryo. The LINC complex, consisting of the KASH protein UNC-83 at the outer nuclear membrane and the SUN protein UNC-84 at the inner nuclear membrane, recruit kinesin-1 to the surface of nuclei through binding to KLC-2, and transmits the forces to inside the nucleus. (B) Quantification of hyp7 nuclear migration. Each point represents the total number of abnormally located (found at the dorsal cord) hyp7 nuclei per animal. n=9 for KLC-2 null. n=14 for hKLC4 G369fs. n=20 for all the other strains. Means with 95% CI are shown in error bars. Unpaired student t-tests were performed on the indicated comparisons; ns means not significant, p>0.05; * p<0.05; **** p<0.0001. (C-F) Lateral view of L1-early L2 (C) *hKLC4*, (D) *hKLC4* T381I), (E) *hKLC4* G369fs, and (F) *hKLC4* G369fs/+ animals expressing hypodermal nuclear GFP. Dashed lines mark the sides of the animal. Dorsal is up. Arrows show abnormally located (in the dorsal cord) hyp7 nuclei. Scale bar, 42.2 µm.

### Functional analysis of *hKLC4* variants of uncertain significance

After showing that the *hKLC4* animals were healthy, our goal was to introduce missense variants into the *hKLC4* avatar to test their possible effects on *C. elegans* development as an indicator of clinical interest. Four *hKLC4* variants were introduced into our *hKLC4* worm line using CRISPR/Cas9 gene editing, chosen as discussed above. Variants *hKLC4* R72H and *hKLC4* R358H were predicted to be begin, and *hKLC4* A295P and *hKLC4* T381I were predicted to be pathogenic (Supplemental Figure 1). We quantified the brood size and lethality of the missense variants and found that neither the predicted benign nor the predicted pathogenic *hKLC4* mutations had a significantly deleterious effect on the percent lethality observed in the parental *hKLC4* line (Figure 2B). However, one of the predicted pathogenic mutations, *hKLC4* T381I, led to a significant decrease in the brood size (Figure 2C).

We had similar results when we observed the swimming behavior of the missense mutants. The two predicted benign variants and the A295P variant had no effect on the swimming rate of the *hKLC4* parental line. However, the predicted disease allele *hKLC4* T381I caused a severe motility defect, with only 0.085 ± 0.038 bbps (Video 3) compared to the parental *hKLC4* worms that had 0.72 ± 0.21 bbps (Figure 3).

In our nuclear migration assay, the benign variants *hKLC4* R72H and R358H did not cause any hyp7 nuclear migration defects (Figure 4). In contrast, both predicted pathogenic variants, *hKLC4* A295P and T381I caused mild, but significant nuclear migration defects where 0.43 ± 0.34 and 0.62 ± 0.24 hyp7 nuclei were observed in the dorsal cord of an average worm (Figure 4B, D).

### Clinical variant *hKLC4* G369fs animals have severe defects

Finally, we introduced the clinical variant of uncertain significance *hKLC4* G369fs using CRISPR/Cas9 genome editing. Attempts to generate a homozygote line failed so we suspected this variant to be lethal. We introduced G369fs into a strain humanized for *hKLC4* that also contained a *klc-2::gfp* rescue array. The rescuing array is expressed from an extrachromosomal array that in *C. elegans* is lost in a high percentage of animals during early embryonic cell divisions (Sakamoto *et al*., 2005). Thus, this strain produces *hKLC4* G369fs animals both with and without the rescuing array. All *hKLC4* G369fs animals that survived to adulthood maintained the *klc-2::gfp* rescuing array, suggesting that all the animals that lost the rescuing array died as embryos or early larvae. We were therefore unable to measure the brood size or swimming ability of the *hKLC4* G369fs animals. However, we were able to observe nuclear migration defects in the rare *hKLC4* G369fs animals that escaped embryonic lethality and could be scored as young larvae before dying. The *hKLC4* G369fs animals had severe nuclear migration defects with a mean of 5.9 ± 1.3 hyp7 nuclei in the dorsal cord of an animal as compared to nearly zero nuclei in the dorsal cord of the *hKLC4* animals (Figure 4). Taken together, these data show that the G369fs mutation is very severe, making the hKLC4 animals very sick.

To test the extent to which the *KLC4* frame-shift variant at residue 369 acts in a haplo-insufficient manner or whether it is the sole contributor to the clinical features, we crossed the truncation mutant to the *hKLC4* strain to assay heterozygotes. Heterozygous *hKLC4* G369fs/+ animals were healthy, viable, and did not have a significant swimming defect (Video 4) compared to *hKLC4* animals (Figure 3A). However, the *hKLC4* G369fs/+ heterozygotes did have a weak, but significant nuclear migration defect in hyp7 precursors (Figure 4B, F). This phenotype was similar to the one observed in predicted pathogenic variants *hKLC4* A295P and *hKLC4* T381I. Thus, while heterozygous *hKLC4* G369fs/+ animals are healthier than the homozygous truncation animals, they still have a significant hyp7 nuclear migration defect.

## Discussion

In this study, we characterized a heterozygous *KLC4* variant of unknown significance detected in an individual with HSP. We engineered and used a humanized *C. elegans* model to test the physiological relevance of the variant in a heterologous *in vivo* system. In conclusion, we have demonstrated that *C. elegans* can be used to model disease-associated variants of human *KLC4*, that we can use the *hKLC4 C. elegans* strain generated here to test the physiological impact of other *KLC4* variants, and that this strategy could be used to model neuromuscular diseases in in other genes with clear orthologs in *C. elegans*, including LINC complexes that target kinesin light chains to the nucleus.

There are thousands of rare diseases, each of which affect fewer than 1/2000 people. About 300-400 million people suffer from rare diseases worldwide (Nguengang Wakap *et al*., 2020). Whole genome sequencing has aided in the diagnosis of rare diseases, but the identification of a genetic underpinning of a disease still fails up to 75% of the time (Smedley *et al*., 2021). We therefore need to develop animal models to test variants of unknown significance to make more efficient and better clinical diagnoses. One goal of the Undiagnosed Disease Network is to bring together clinicians and basic scientists to test variants of unknown significance in animal models (Baldridge *et al*., 2021). Here, we report clinical findings implicating a novel variant in the kinesin light chain gene *KLC4* from a proband with HSP and the subsequent generation of a humanized *C. elegans* model to test the significance of the variant.

We identified an individual with late-onset HSP with a heterozygous variant in *KLC4* predicted to cause a frame shift at residue 369, closely followed by a premature stop codon. An additional family was previously reported where a premature stop codon after residue 277 of *KLC4* caused HSP in a recessive manner; heterozygous family members did not have any symptoms (Bayrakli *et al*., 2015). Truncations in *KLC4* after either 277 or 369 residues are predicted to disrupt the TPR domain, which mediates the interaction between kinesin and the cargo adaptor (Pernigo *et al*., 2013; Zhu *et al*., 2012), suggesting that both *KLC4* variants should produce similar pathologies. They could be acting in a dominant-negative manner in the proband, which would not be recapitulated in *C. elegans* due to nonsense-mediated decay of the message. Alternatively, there could be a second variant in the proband acting synergistically with the *KLC4* truncation. Variants in other genes were identified by whole genome sequencing of the individual and ongoing studies are attempting to determine their contributions to pathogenicity. If such a variant were identified, it could lead to significant insights in the function of *KLC4* in normal development and the progression of HSP.

We aimed to make a humanized *C. elegans* model to test clinical *KLC4* variants. The open reading frame of the *C. elegans* ortholog *klc-2* was successfully replaced with the human *KLC4* coding sequence. While the *klc-2* null alleles are not viable(Sakamoto *et al*., 2005), the *hKLC4* model had only low levels of lethality and a slightly reduced brood size. However, the *hKLC4* animals had significant swimming defects. Thus, the humanized model expressing *KLC4* under control of the endogenous *klc-2* locus retained much, but not all of the function of *klc-2*. We used these phenotypes as a baseline to compare with the effects caused by the introduction of variants of unknown significance into the *hKLC4* line, including the clinical truncation variant. The clinical variant was homozygous lethal, and escaper larvae had severe nuclear migration defects like those with *klc-2* null alleles (Fridolfsson *et al*., 2010), suggesting that the clinical variant would lead to severe pathologies when homozygous. We next examined whether the *KLC4* frame-shift variant at residue 369 acts in a haplo-insufficient manner. We observed a mild nuclear migration defect in heterozygous animals, consistent with other disease-associated variants. This mild phenotype in *C. elegans* could therefore be useful in predicting whether a variant of unknown significance might cause clinical symptoms of HSP. Further work is needed to determine if a heterozygous loss-of-function *KLC4* variant can cause HSP; this includes finding more affected individuals who harbor disease-causing *KLC4* variants.

We next used the *hKLC4 C. elegans* line to test predicted homozygous variants of unknown significance with single amino acid changes that are predicted to be disruptive and potentially pathogenic to *hKLC4*. One of the four variants, *hKLC4* T381I, disrupted the function of *hKLC4* in *C. elegans*. The T381I variant had significant phenotypes including a reduced brood size, slower thrashing in our swimming assay, and an increased number of hyp7 precursor nuclei that failed to migrate. These data suggest that the T381I variant is also likely to disrupt KLC4 function in human cells, while the other three tested variants are unlikely to be pathogenic. These experiments demonstrate that the *hKLC4 C. elegans* strain produced here is a useful tool to test the potential pathogenicity of variants of unknown significance in *KLC4* found in future clinical cases.

The kinesin light chain is part of a network of proteins conserved from *C. elegans* to humans including the LINC complex-forming KASH and SUN proteins (Starr, 2019; Fridolfsson and Starr, 2010; Roux *et al*., 2009). Variants in the genes encoding components of human LINC complexes, including Nesprins, have been implicated in a wide variety of diseases, including neurological disorders, muscular dystrophies, and various cancers (Janin *et al*., 2017). Humanized *C. elegans* strains for LINC complex components would provide additional reagents to test clinical variants in this important complex. The success of the humanized *KLC4 C. elegans* line described here suggests that this approach may be feasible for modeling other neuromuscular diseases associated with LINC complex dysfunction.

## Acknowledgements

We thank members of the Starr-Luxton lab for their helpful comments, especially Daniel Elnatan for help with the AlphaFold figure. This work was supported by the NIH Common Fund, through the Office of Strategic Coordination/Office of the NIH Director under Award Numbers U01HG007530 and U01HG007672. The content is solely the responsibility of the authors and does not necessarily represent the official views of the National Institutes of Health. This research was also supported by NIH grant R35GM134859 to DAS.

---

**Video 1:**
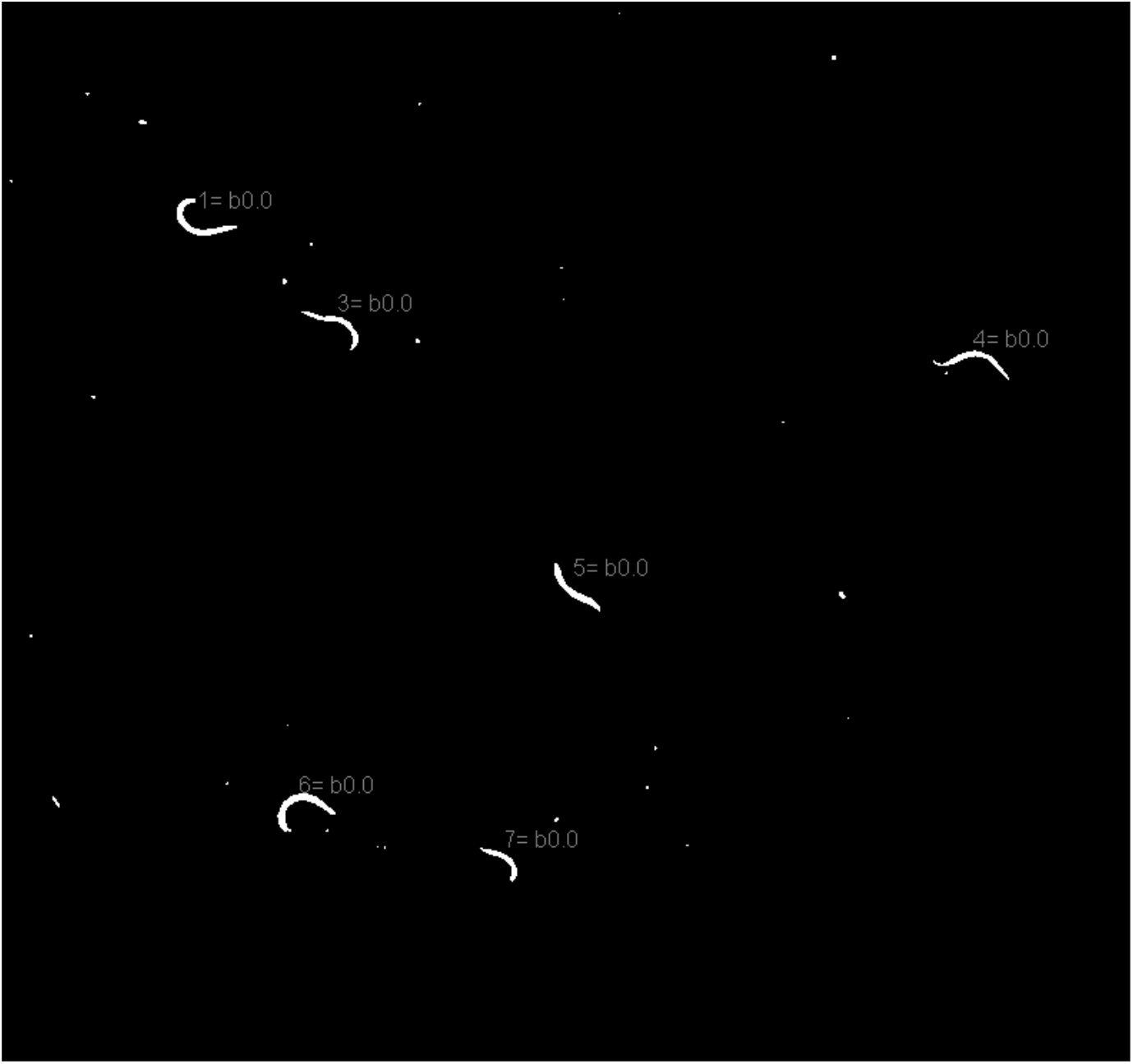
Wildtype (N2) worms have no motility defects. L4-stage worms swimming in buffer. Fiji wrMTrck plugin (Nussbaum-Krammer *et al*., 2015) was used to track each worm to measure the number of body bends per second (BBPS).

**Video 2:**
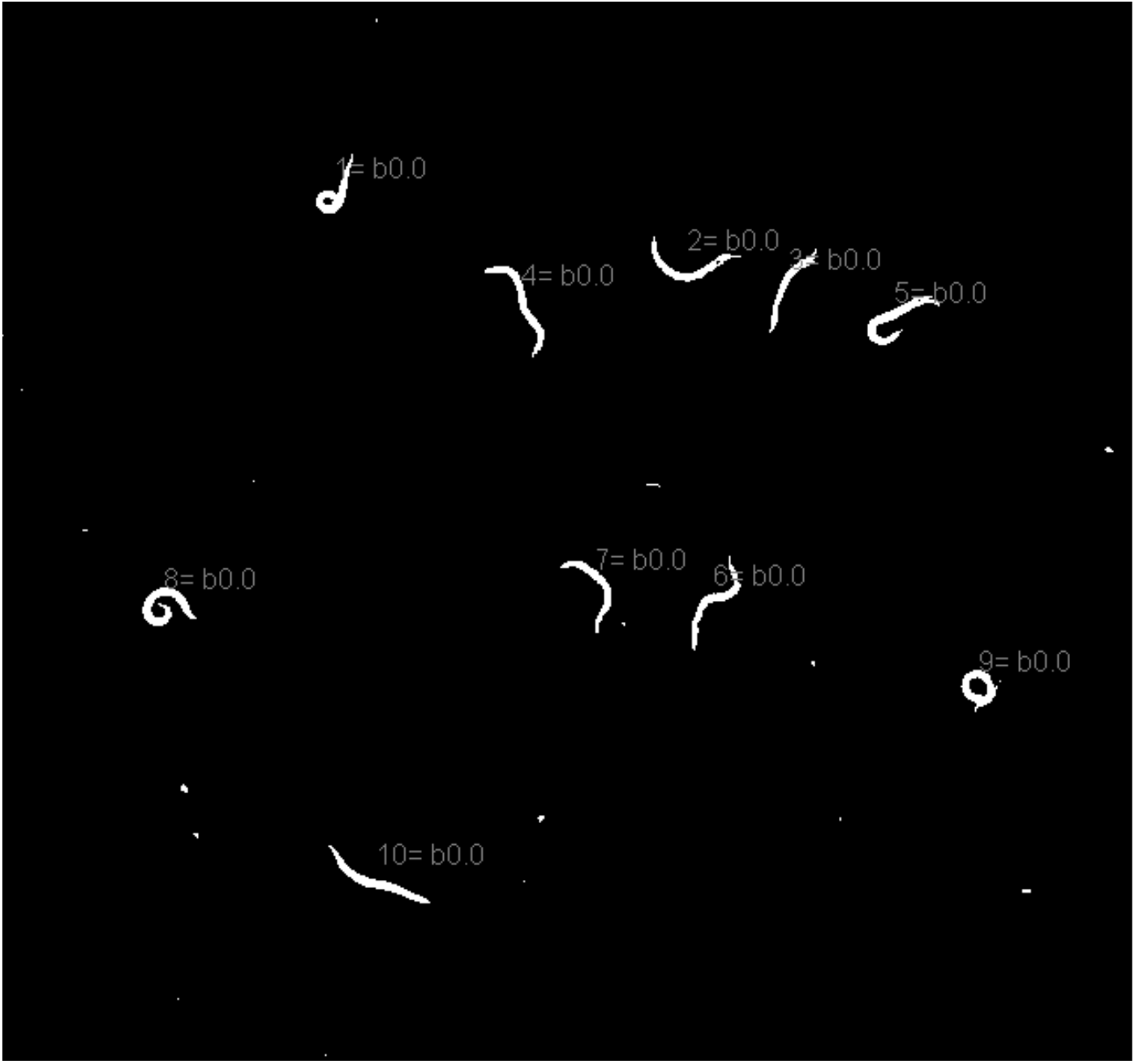
*hKLC4* worms have a significant motility defects, but they retain most of their swimming ability. L4-stage worms swimming in buffer. Fiji wrMTrck plugin(Nussbaum-Krammer *et al*., 2015) was used to track each worm to measure the number of body bends per second (BBPS).

**Video 3:**
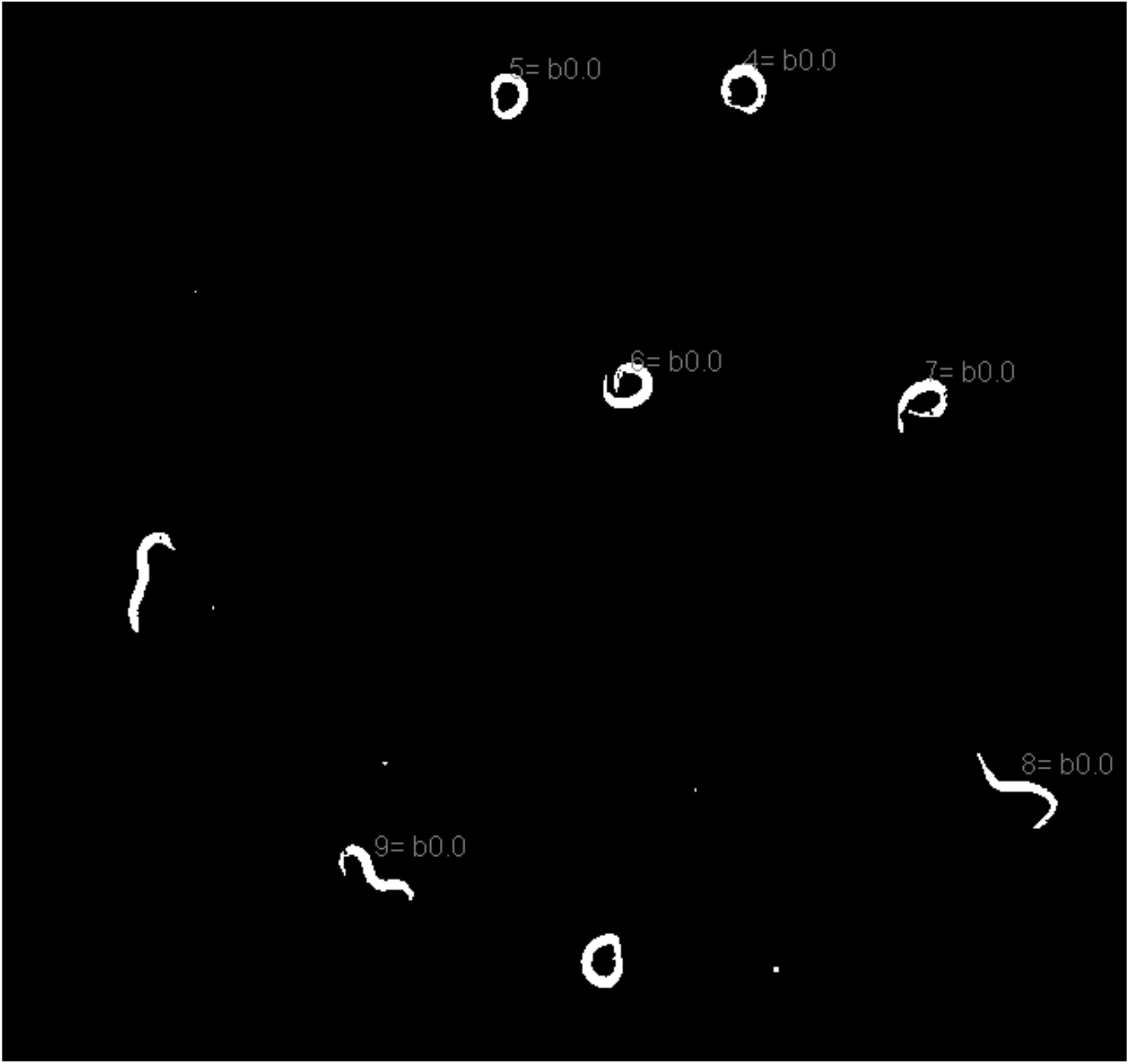
*hKLC4* T381I worms have a severe motility defect. L4-stage worms swimming in buffer. Fiji wrMTrck plugin(Nussbaum-Krammer *et al*., 2015) was used to track each worm to measure the number of body bends per second (BBPS).

**Video 4:**
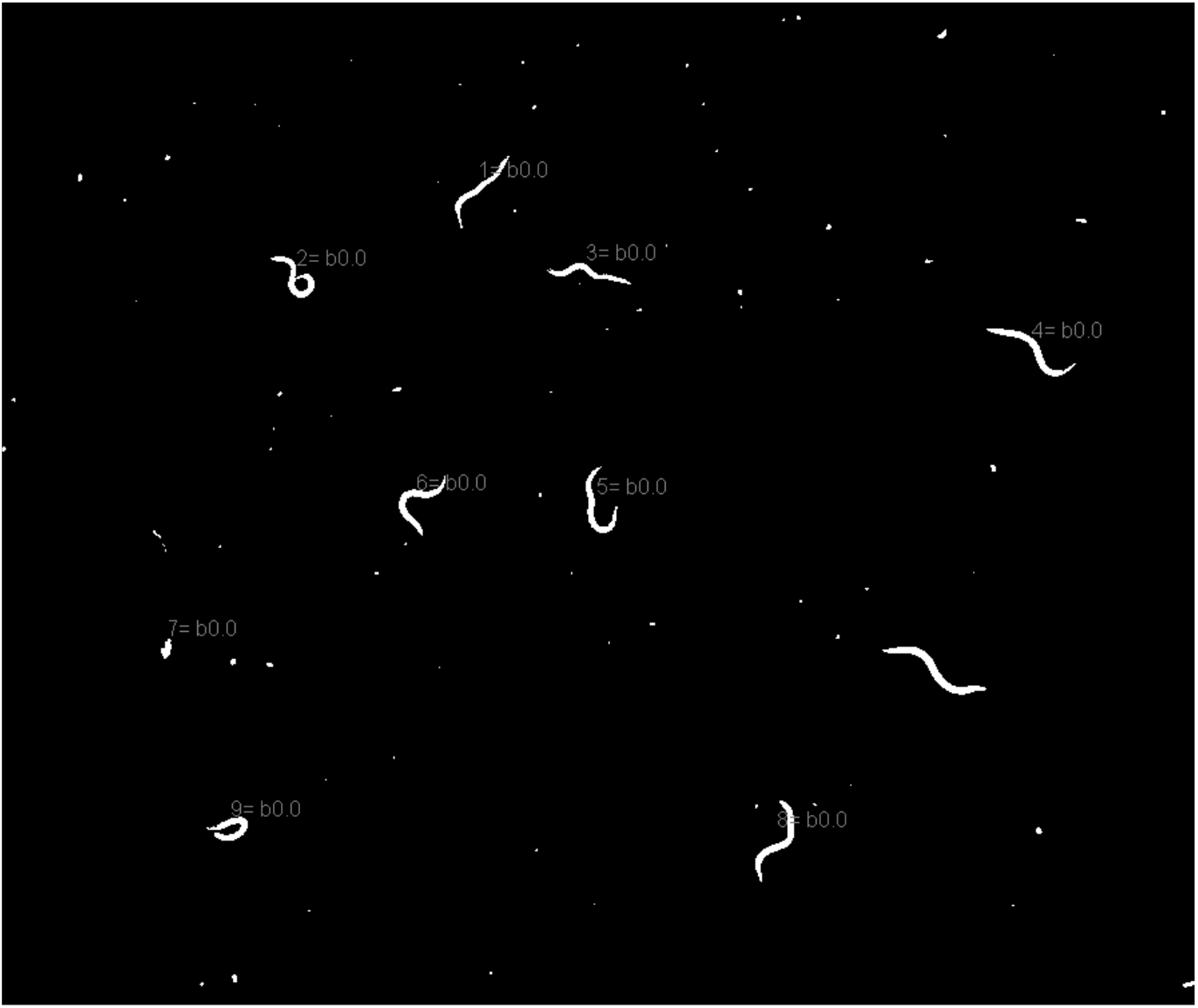
*hKLC4* G369fs/+ worms have a significant motility defect, but they retain most of their swimming ability. L4-stage worms swimming in buffer. Fiji wrMTrck plugin(Nussbaum-Krammer *et al*., 2015) was used to track each worm to measure the number of body bends per second (BBPS).

**Supplemental Figure 1:**
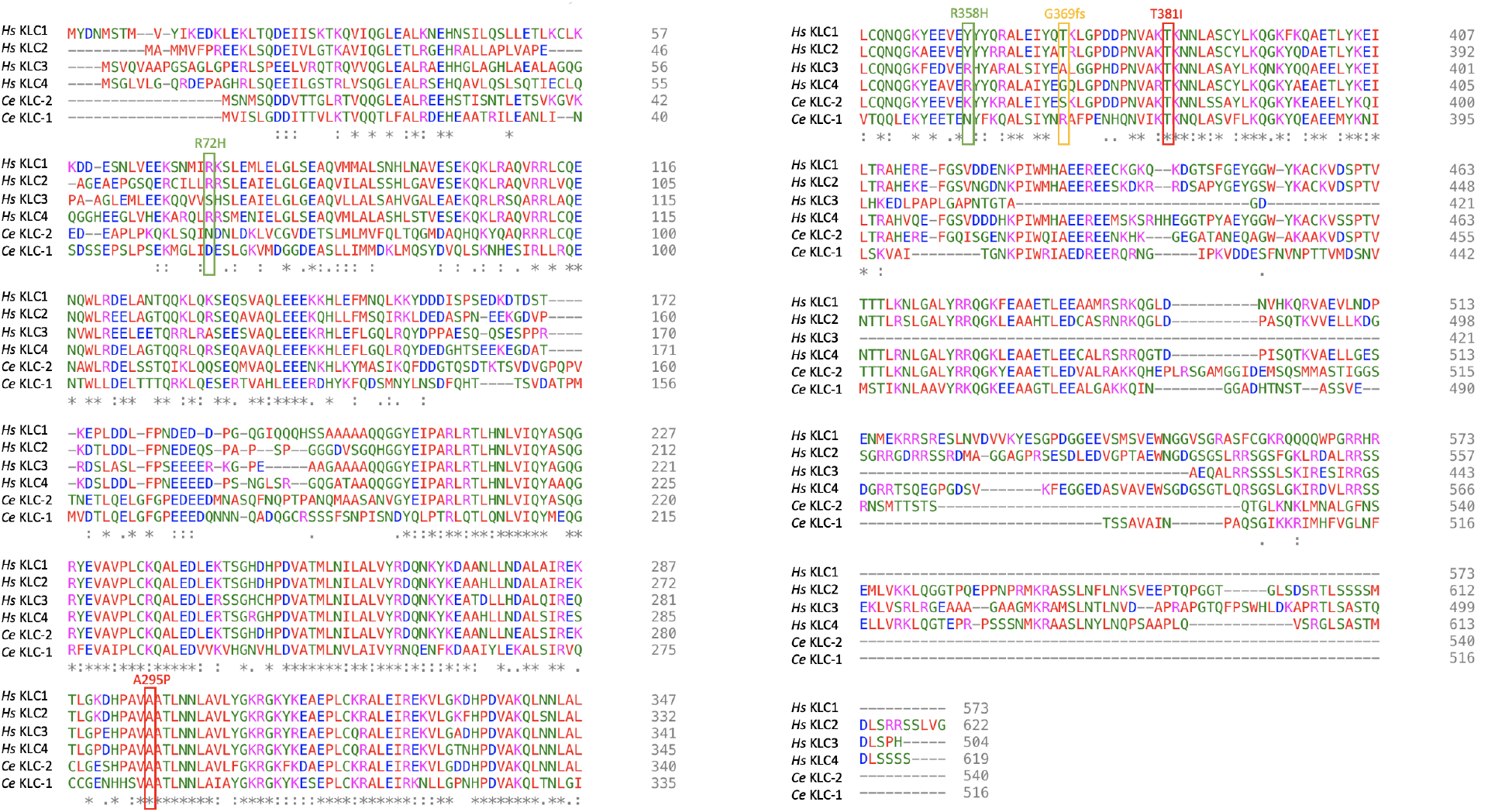
Alignment of human and *C. elegans* kinesin light chains. A protein sequence alignment of the four human and two *C. elegans* kinesin light chains are shown. Amino acids are color coded based on side chain properties. The five residues introduced into the *hKLC4* line of *C. elegans* are boxed. R72H and R358H (green boxes) missense mutations are predicted to be benign. A295P and T381I (red boxes) missense mutations predicted to be pathogenic. G369fs (orange box) is the clinical variant of uncertain significance that introduces a frame shift and early stop. The notations underneath the residues show how conserved each residue is;. indicates at least 50% identity, : indicates a higher level of identity/similarity, and * indicates that the residue is identical in all 6 kinesin light chains.

**Supplemental Figure 2:**
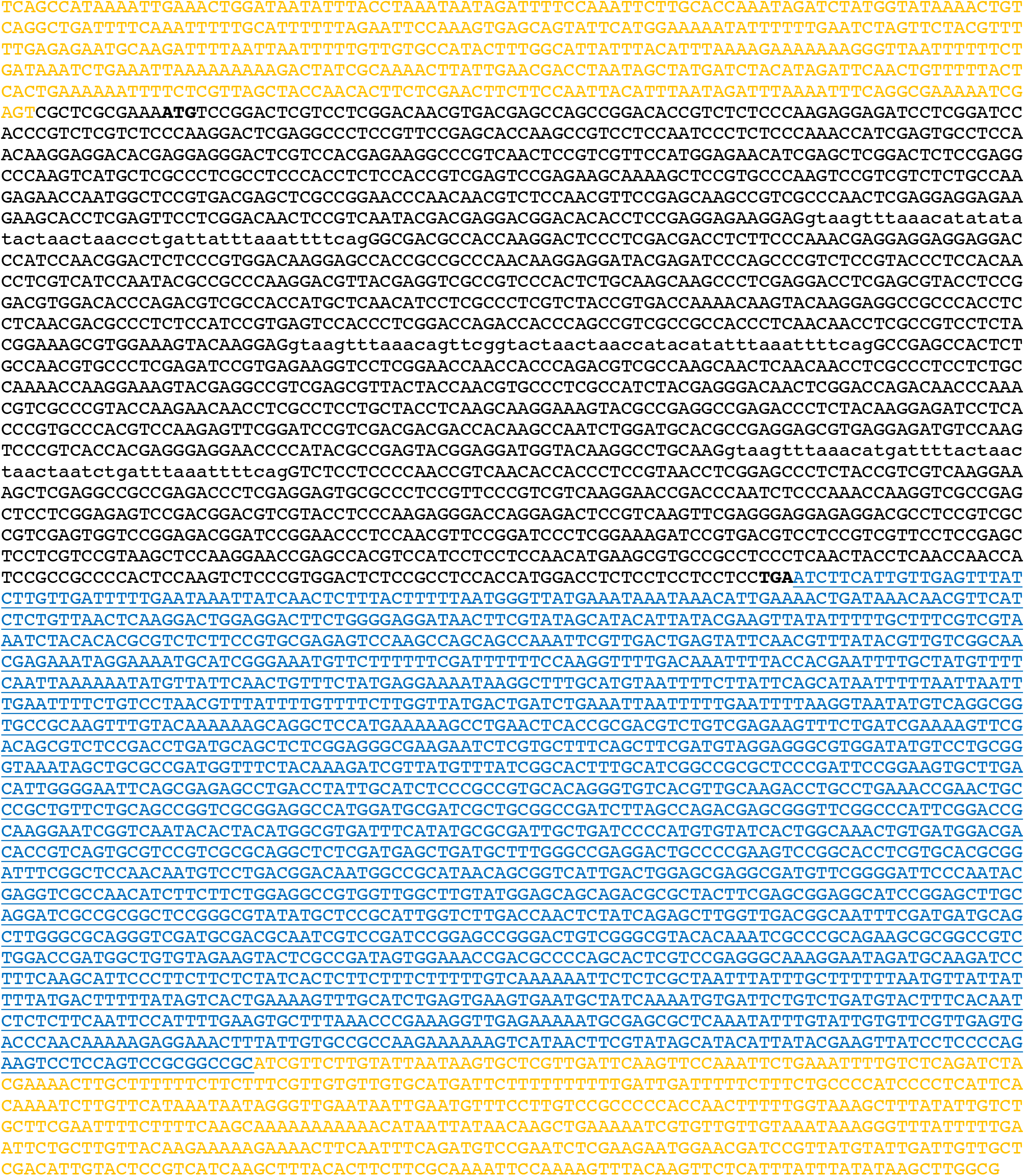
The *hKLC4* sequence and hygromycin resistance selection cassette inserted into the *klc-2* locus. Homology arms that are the native sequence of upstream and downstream of the *klc-2* open reading frame are shown in yellow. Codon-optimized hKLC4 sequence is shown in black. Start and stop codons are bolded. Exons are shown in capital letters and synthetic introns are shown in lower-case letters. The hygromycin gene resistance selection cassette is shown in blue.

## References

AdzhubeIvan, I. A., Schmidt, S., Peshkin, L., Ramensky, V. E., Gerasimova, A., Bork, P., Kondrashov, A. S., Sunyaev, S. R. (2010). A Method and Server for Predicting Damaging Missense Mutations. Nat Methods. 7 (4): 248–249. https://doi.org/10.1016/0002-9343(63)90102-5.

Arribere, J. A., Bell, R.T., Fu, B.X.H., Artiles, K.L., Hartman, P.S., Fire, A.Z. (2014). Efficient Marker-Free Recovery of Custom Genetic Modifications with CRISPR/Cas9 in Caenorhabditis Elegans. Genetics. 198 (3): 837–46. https://doi.org/10.1534/genetics.114.169730.

Baek, J-H., Lee J., Yun H.S., Lee C-W., Song J-Y., Um, H.D., Park, J.K. et al. (2018). Kinesin Light Chain-4 Depletion Induces Apoptosis of Radioresistant Cancer Cells by Mitochondrial Dysfunction via Calcium Ion Influx Article. Cell Death and Disease. 9, 496. https://doi.org/10.1038/s41419-018-0549-2.

Baek, J-H., Yun H.S., Kim, J.Y., Lee, J., Lee, Y.J., Lee, C-W., Song, J.Y. et al. (2020). Kinesin Light Chain 4 as a New Target for Lung Cancer Chemoresistance via Targeted Inhibition of Checkpoint Kinases in the DNA Repair Network. Cell Death and Disease. 11, 398. https://doi.org/10.1038/s41419-020-2592-z.

Baldridge, D., Wangler, M.F., Bowman, A.N., Yamamoto, S., Schedl, T., Pak, S.C., Postlethwait, J.H. et al. (2021). Model Organisms Contribute to Diagnosis and Discovery in the Undiagnosed Diseases Network: Current State and a Future Vision. Orphanet Journal of Rare Diseases. 16 (1): 1–17. https://doi.org/10.1186/s13023-021-01839-9.

Bayrakli, F., Poyrazoglu, H.G., Yuksel, S., Yakicier, C., Erguner, B., Sagiroglu, M.S., Yuceturk, B. et al. (2015). Hereditary Spastic Paraplegia with Recessive Trait Caused by Mutation in KLC4 Gene. J Hum Gen. 60 (12): 763–68. https://doi.org/10.1038/jhg.2015.109.

Blumenthal, T. (2012). Trans-Splicing and Operons in C. Elegans. In Wormbook Online Rev C Elegans Biology, 1–11.

Bone, C.R., Tapley, E.C., Gorjánácz, M., and Starr, D.A. (2014). The Caenorhabditis Elegans SUN Protein UNC-84 Interacts with Lamin to Transfer Forces from the Cytoplasm to the Nucleoskeleton during Nuclear Migration. Molecular Biology of the Cell 25 (18): 2853–65. https://doi.org/10.1091/mbc.E14-05-0971.

Brenner, S. (1974). The Genetics of Caenorhabditis Elegans. Genetics. 77 (1): 71–94. https://doi.org/10.1080/01677063.2020.1802723.

Chen, C., Fenk, L.A., and De Bono, M. (2013). Efficient Genome Editing in Caenorhabditis Elegans by CRISPR-Targeted Homologous Recombination. Nucleic Acids Res. 41 (20). https://doi.org/10.1093/nar/gkt805.

Farboud, B., Severson, A. F., Meyer, B.J. (2019). Strategies for Efficient Genome Editing Using CRISPR-Cas9. Genetics. 211 (February): 431–57.

Ferreira, C.R. (2019). The Burden of Rare Diseases. Am J Med Genet A. 179 (6): 885–92. https://doi.org/10.1002/ajmg.a.61124.

Fridolfsson, H.N., Ly, N., Meyerzon, M., and Starr, D.A. (2010). UNC-83 Coordinates Kinesin-1 and Dynein Activities at the Nuclear Envelope during Nuclear Migration. Dev Biol. 338 (2): 237–50. https://doi.org/10.1016/j.ydbio.2009.12.004.

Fridolfsson, H.N., and Starr, D.A. (2010). Kinesin-1 and Dynein at the Nuclear Envelope Mediate the Bidirectional Migrations of Nuclei. J Cell Biol. 191 (1): 115–28. https://doi.org/10.1083/jcb.201004118.

Fridolfsson, H.N., Herrera, L.A., Brandt, J.N., Cain, N.E., Hermann, G.J., and Starr, D.A. (2018). Genetic Analysis of Nuclear Migration and Anchorage to Study LINC Complexes During Development of Caenorhabditis Elegans. Methods Mol Biol. 1840, 163–180. doi:10.1007/978-1-4939-8691-0_13.

Giudice, M. L., Neri, M., Falco, M., Sturnio, M., Calzolari, E., Di Benedetto, D., and Fichera, M. (2014). A Missense Mutation in the Coiled-Coil Domain of the KIF5A Gene and Late-Onset Hereditary Spastic Paraplegia. Arch Neurol. 63 (2): 284–87. doi:10.1001/archneur.63.2.284

Gumeni, S., Vantaggiato, C., Montopoli, M. and Orso, G. (2021). Hereditary Spastic Paraplegia and Future Therapeutic Directions: Beneficial Effects of Small Compounds Acting on Cellular Stress. Front Neuro. 15 (May): 1–16. https://doi.org/10.3389/fnins.2021.660714.

Guo, X., Zhang, T., Hu, Z., Zhang, Y., Shi, Z., Wang, Q., Cui, Y., Wang, F., Zhao, H. and Chen, Y. (2014). Efficient RNA/Cas9-Mediated Genome Editing in Xenopus Tropicalis. Development (Cambridge). 141 (3): 707–14. https://doi.org/10.1242/dev.099853.

Hirokawa, N., Niwa, S. and Tanaka, Y. (2010). Molecular Motors in Neurons: Transport Mechanisms and Roles in Brain Function, Development, and Disease. Neuron. 68 (4): 610–38. https://doi.org/10.1016/j.neuron.2010.09.039.

Hopkins, C.E., Brock, T., Caulfield, T.R. and Bainbridge, M. (2022). Phenotypic Screening Models for Rapid Diagnosis of Genetic Variants and Discovery of Personalized Therapeutics. Mol Aspects Med. https://doi.org/10.1016/j.mam.2022.101153.

Ioannidis, N.M., Rothstein, J.H., Pejaver, V., Middha, S., McDonnell, S.K., Baheti, S., Musolf, A., et al. (2016). REVEL: An Ensemble Method for Predicting the Pathogenicity of Rare Missense Variants. Am J Hum Genet. 99 (4): 877–85. https://doi.org/10.1016/j.ajhg.2016.08.016.

Janin, A., Bauer, D., Ratti, F., Millat, G. and Méjat, A. (2017). Nuclear Envelopathies: A Complex LINC between Nuclear Envelope and Pathology. Orphanet J Rare Diseases. 12 (1): 1–16. https://doi.org/10.1186/s13023-017-0698-x.

Jumper, J., Evans, R., Pritzel, A., Green, T., Figurnov, M., Ronneberger, O., Tunyasuvunakool, K., Bates, R., Zidek, A., Potapenko, A., et al. (2021). Highly Accurate Protein Structure Prediction with AlphaFold. Nature. 596 (7873): 583–89. https://doi.org/10.1038/s41586-021-03819-2.

Junco, A., Bhullar, B., Tarnasky, H. A. and Van der Hoorn, F. A. (2001). Kinesin Light-Chain KLC3 Expression in Testis Is Restricted to Spermatids. Biol Rep. 64 (5): 1320–30. https://doi.org/10.1095/biolreprod64.5.1320.

Karczewski, K. J., Francioli, L. C., Tiao, G., Cummings, B. B., JAlföldi, J., Wang, Q., Collins, R. L., Laricchia, K. M., Ganna, A., Birnbaum, D. P. et al. (2020). The Mutational Constraint Spectrum Quantified from Variation in 141,456 Humans. Nature. 581 (7809): 434–43. https://doi.org/10.1038/s41586-020-2308-7.

Kircher, M., Witten, D.M., Jain, P., O’Roak, B.J., Cooper, G.M. and Shendure, J. (2014). A General Framework for Estimating the Relative Pathogenicity of Human Genetic Variants. Nat Genet. 46 (3): 310–15. https://doi.org/10.1038/ng.2892.

Kropp, P. A., Bauer, R., Zafra, I., Graham, C., and Golden, A. (2021). Caenorhabditis Elegans for Rare Disease Modeling and Drug Discovery: Strategies and Strengths.” DMM. 14 (8). https://doi.org/10.1242/DMM.049010.

Kurd, D. D., and Saxton, W. M. (1996). Kinesin Mutations Cause Motor Neuron Disease Phenotypes by Disrupting Fast Axonal Transport in Drosophila. Genetics. 144 (3): 1075– 85. https://doi.org/10.1093/genetics/144.3.1075.

Mandelkow, E., and Mandelkow, E-M. (2002). Kinesin Motors and Disease. Trends in Cell Biol. 12 (12): 585–91. https://doi.org/10.1016/s0962-8924(02)02400-5

Mattout, A., Pike, B. L., Towbin, B. D., Bank, E. M., Gonzalez-Sandoval, A., Stadler, M. B., Meister, P., Gruenbaum, Y., and Gasser, S. M. (2011). An EDMD Mutation in C. Elegans Lamin Blocks Muscle-Specific Gene Relocation and Compromises Muscle Integrity. Curr Biol. 21 (19): 1603–14. https://doi.org/10.1016/j.cub.2011.08.030.

Meyerzon, M., Fridolfsson, H. N., Ly, N., McNally, F. J. and Starr, D. A. (2009). UNC-83 Is a Nuclear-Specific Cargo Adaptor for Kinesin-1-Mediated Nuclear Migration. Development. 136 (16): 2725–33. https://doi.org/10.1242/dev.038596.

Mitreva, M., Wendl, M. C., Martin, J., Wylie, T., Yin, Y., Larson, A., Parkinson, J., Waterston, R. H. and McCarter, J. P. (2006). Codon Usage Patterns in Nematoda: Analysis Based on over 25 Million Codons in Thirty-Two Species. Genome Biol. 7 (8). https://doi.org/10.1186/gb-2006-7-8-r75.

Nguengang Wakap, S., Lambert, D. M., Olry, A., Rodwell, C., Gueydan, C., Lanneau, V., Murphy, D., Le Cam, Y. and Rath, A. (2020). Estimating Cumulative Point Prevalence of Rare Diseases: Analysis of the Orphanet Database. Eur J Hum Genet. 28 (2): 165–73. https://doi.org/10.1038/s41431-019-0508-0.

Nussbaum-Krammer, C. I., Mário, N. F., Brielmann, R. M., Pedersen, J. S. and Morimoto, R. I. (2015). Investigating the Spreading and Toxicity of Prion-like Proteins Using the Metazoan Model Organism C. Elegans. J Vis Exp. 52321 (95): 1–15. https://doi.org/10.3791/52321.

Paix, A., Folkmann, A., Rasoloson, D. and Seydoux, G. (2015). High Efficiency, Homology-Directed Genome Editing in Caenorhabditis Elegans Using CRISPR-Cas9 Ribonucleoprotein Complexes. Genetics 201 (1): 47–54. https://doi.org/10.1534/genetics.115.179382.

Parodi, L., Fenu, S., Stevanin, G. and Durr, A. (2017). Hereditary Spastic Paraplegia: More than an Upper Motor Neuron Disease. Revue Neurologique 173 (5): 352–60. https://doi.org/10.1016/j.neurol.2017.03.034.

Perlson, E., Maday, S., Fu, M-M., Moughamian, A. J. and Holzbaur, E. L. (2010). Retrograde Axonal Transport: Pathways to Cell Death? Trends Neuorosci 33 (7): 335–344. https://doi.org/10.1016/j.tins.2010.03.006.

Pernigo, S., Lamprecht, A., Steiner, R. A. and Dodding, M. P. (2013). Structural Basis For Kinesin-1:Cargo Recognition. Science 340 (6130): 356–59. https://doi.org/10.1126/science.1234264.

Pierce-Shimomura, J. T., Chen, B. L., Mun, J. J., Ho, R., Sarkis, R. and McIntire, S. L. (2008). Genetic Analysis of Crawling and Swimming Locomotory Patterns in C. Elegans. Proc Natl Acad Sci U S A. 105 (52): 20982–87. https://doi.org/10.1073/pnas.0810359105.

Rahman, A., Friedman, D. S. and Goldstein, L. S. B. (1998). Two Kinesin Light Chain Genes in Mice. J Biol Chem. 273 (25): 15395–403. https://doi.org/10.1074/jbc.273.25.15395.

Ross, J. L., Ali, Y. M. and Warshaw, D. M. (2008). Cargo Transport: Molecular Motors Navigate a Complex Cytoskeleton. Curr Opin Cell Biol. 20 (1): 41–47. https://doi.org/10.1016/j.ceb.2007.11.006.C

Roux, K. J., Crisp, M. L., Liu, Q., Kim, D., Kozlov, S., Stewart, C. L. and Burke, B. (2009). Nesprin 4 Is an Outer Nuclear Membrane Protein that Can Induce Kinesin-Mediated Cell Polarization. Proc Natl Acad Sci U S A. 106 (7): 2194–99. https://doi.org/10.1073/pnas.0808602106.

Sakamoto, R., Byrd, D. T., Brown, H. M., Hisamoto, N., Matsumoto, K. and Jin, Y. (2005). The Caenorhabditis Elegans UNC-14 RUN Domain Protein Binds to the Kinesin-1 and UNC-16 Complex and Regulates Synaptic Vesicle Localization. Mol Biol Cell. 16 (2): 483– 96. https://doi.org/10.1091/mbc.E04-07-0553.

Saxton, W. M. and Hollenbeck, P.J. (2012). The Axonal Transport of Mitochondria. J Cell Sci. 125 (9): 2095–2104. https://doi.org/10.1242/jcs.053850.

Shribman, S., Reid, E., Crosby, A. H., Houlden, H. and Warner, T. T. (2019). Hereditary Spastic Paraplegia: From Diagnosis to Emerging Therapeutic Approaches. Lancet Neurol. 18 (12): 1136–46. https://doi.org/10.1016/S1474-4422(19)30235-2.

Sim, N. L., Kumar, P., Hu, J., Henikoff, S., Schneider, G. and Ng, P. C. (2012). SIFT Web Server: Predicting Effects of Amino Acid Substitutions on Proteins. Nucleic Acids Res. 40 (W1): 452–57. https://doi.org/10.1093/nar/gks539.

Smedley, D., Smith, K. R., Martin, A., Thomas, E. A., McDonagh, E. M., Cipriani, V., Ellingford, J. M. et al. (2021). 100,000 Genomes Pilot on Rare-Disease Diagnosis in Health Care — Preliminary Report. N Engl J Med. 385 (20): 1868–80. https://doi.org/10.1056/nejmoa2035790.

Starr, D. A. (2019). A Network of Nuclear Envelope Proteins and Cytoskeletal Force Generators Mediates Movements of and within Nuclei throughout Caenorhabditis Elegans Development.” Exp Biol Med. 244 (15): 1323–32. https://doi.org/10.1177/1535370219871965.

Sulston, J. E., Schierenberg, E., White, J. G. and Thomson, J. N. (1983). The Embryonic Cell Lineage of the Nematode Caenorhabditis Elegans. Dev Biol. 100 (1): 64–119. https://doi.org/10.1016/0012-1606(83)90201-4.

Taiber, S., Gozlan, O., Cohen, R., Andrade, L. R., Gregory, E. F., Starr, D. A., Moran, Y., Hipp R., Kelley, M. W., Manor, U. et al. (2022). A Nesprin-4/Kinesin-1 Cargo Model for Nuclear Positioning in Cochlear Outer Hair Cells. Front Cell Dev Biol. 10 (September): 1– 15. https://doi.org/10.3389/fcell.2022.974168.

Vale, R. D., Reese, T. S. and Sheetz, M. P. (1985). Identification of a Novel Force-Generating Protein, Kinesin, Involved in Microtubule-Based Motility. Cell 42 (1): 39–50. https://doi.org/10.1016/S0092-8674(85)80099-4

Verhey, K. J., Lizotte, D. L., Abramson, T., Barenboim, L., Schnapp, B. J. and Rapoport, T. A. (1998.) Light Chain-Dependent Regulation of Kinesin’s Interaction with Microtubules J Cell Biol. 143 (4): 1053–66. https://doi.org/10.1083/jcb.143.4.1053.

Yates, A., Akanni, W., Amode, M. R., Barrell, D., Billis, K., Carvalho-Silva, D., Cummins, C. et al. (2016). Ensembl 2016. Nucleic Acids Res. 44 (D1): D710–16. https://doi.org/10.1093/nar/gkv1157.

Zhang, Y., Ou, Y., Cheng, M., Shojaei Saadi, H., Thundathil, J. C. and van der Hoorn, F. A. (2012). KLC3 Is Involved in Sperm Tail Midpiece Formation and Sperm Function. Dev Biol. 366 (2): 101–10. https://doi.org/10.1016/j.ydbio.2012.04.026.

Zhu, H., Lee, H. Y., Tong, Y., Hong, B. S., Kim, K. P., Shen, Y., Lim, K. J., Mackenzie, F., Tempel, W. and Park, H. W. (2012). Crystal Structures of the Tetratricopeptide Repeat Domains of Kinesin Light Chains: Insight into Cargo Recognition Mechanisms. PLoS ONE 7 (3): 1–10. https://doi.org/10.1371/journal.pone.0033943.

